# Computational modelling reveals neurobiological contributions to static and dynamic functional connectivity patterns

**DOI:** 10.1101/2024.10.01.614888

**Authors:** Linnea Hoheisel, Hannah Hacker, Gereon R Fink, Silvia Daun, Joseph Kambeitz

## Abstract

Functional connectivity (FC) is a widely used indicator of brain function in health and disease, yet its neurobiological underpinnings still need to be firmly established. Recent advances in computational modelling allow us to investigate the relationship between FC and neurobiology non-invasively. These techniques allow for targeted manipulations to study the effect of network disturbances on FC. Most modelling research has concentrated on replicating empirical static FC (sFC). However, FC changes over time, and its dynamic properties are closely linked to behaviour and symptomatology.

In this study, we adapted computational models to reflect both sFC and dynamic FC (dFC) of individuals, allowing for a more comprehensive characterisation of the neurobiological origins of FC. We modelled the brain activity of 200 healthy individuals based on empirical resting-state functional magnetic resonance imaging (fMRI) and diffusion tensor imaging (DTI) data. Simulations were conducted using a group-averaged structural connectome and four parameters guiding regional brain activity: i) G, a global coupling scaling parameter; ii) J_i, the local inhibitory current; iii) J_NMDA, the excitatory NMDA synaptic coupling parameter; and iv) w_p, the excitatory population recurrence weight. We evaluated the models based on four metrics: a) the sFC, b) the FC variance, c) the temporal correlation (TC), and d) the node cohesion (NC). The optimal model for each subject was identified by the fit to both sFC and TC. We analysed associations between brain-wide sFC and TC features with optimal model parameters and fits with a univariate correlation approach and multivariate prediction models. In addition, we used a group-average perturbation approach to investigate the effect of coupling in each region on overall network connectivity.

Our models could replicate empirical sFC and TC but not the FC variance and NC. Both fits and parameters exhibited strong associations with brain connectivity. G correlated positively and J_NMDA negatively with a range of static and dynamic FC features (|r| > 0.2, p(FDR) < 0.05). TC fit correlated negatively, and sFC fit positively with static and dynamic FC features. TC features were predictive of TC fit, sFC features of sFC fit (R^2^ > 0.5). Perturbation analysis revealed that the sFC fit was most impacted by coupling changes in the left paracentral gyrus (Δr = 0.07). In contrast, the left pars triangularis impacted the TC fit most strongly (Δr = 0.24).

Our findings indicate that neurobiological characteristics are associated with individual variability in sFC and dFC, and that sFC and dFC are shaped by small sets of distinct regions. In addition, we show that brain network modelling can replicate some, but not all, properties of dFC, and model fits are strongly influenced by specific FC patterns. By modelling both sFC and dFC, we could produce new insights into neurobiological mechanisms of brain network configurations.

## Introduction

Resting-state functional connectivity (FC), defined as the correlation of activity between different brain regions at rest, is a key signature of brain functioning (Finn et al., 2015; Ooi et al., 2022; van den Heuvel & Hulshoff Pol, 2010). It is related to cognitive ability and personality (Kong et al., 2019; Sripada et al., 2020) as well as the brain’s health status and individual symptoms (Diener et al., 2012; Gallo et al., 2023; Mehta et al., 2021; Parkes et al., 2020; D. Wang et al., 2020). Recent research has indicated that FC is not static but changes over time, exhibiting dynamic characteristics related to behaviour and clinical features (Hutchison et al., 2013; Patanaik et al., 2018; Hoheisel et al., 2024). However, the biological processes underlying both static and dynamic FC remain unclear. Novel computational approaches allow us to investigate these mechanisms by simulating brain activity using empirical neuroimaging data (H. E. Wang et al., 2024). As previous models primarily focussed on replicating static FC, they were not suitable for the investigation of dynamic aspects of FC, which are crucial for understanding brain connectivity and related disorders.

Individual brain FC patterns are influenced by several mechanisms. The role of structural connectivity (SC), which represents the pattern of anatomical links between brain regions, in static FC origin has long been established (Honey et al., 2009; Sporns et al., 2000). However, SC does not account for the entirety of static FC variability between subjects, and cannot account for patterns of dynamic changes in FC (Liégeois et al., 2020). Mechanisms of neurotransmission and neuromodulation also regulate communication between regions (Brink et al., 2016; Hansen, Shafiei, et al., 2022) through factors such as dopaminergic and serotonergic signalling (Klaassens et al., 2015), and neuroreceptor expression patterns (Hansen et al., 2022). In addition, neurobiological characteristics of brain areas which determine regional activity, especially the balance between excitatory and inhibitory populations, impact FC (Gu et al., 2019; Kapogiannis et al., 2013; Levar et al., 2019). In line with this, disruptions of *N*-methyl-D-aspartate (NMDA)-related excitatory signalling have been suggested as a possible mechanism leading to increased connectivity across the brain (Anticevic et al., 2015; Driesen et al., 2013) and increased network flexibility in dynamic FC (Braun et al., 2016).

Brain network modelling allows for a holistic analysis of SC and FC by inferring neurobiological processes from data acquired via magnetic resonance imaging (MRI) (Breakspear, 2017; Schirner et al., 2018; Shine et al., 2021). These models describe regional brain activity using a set of equations with variable parameters governing excitation-inhibition balance and the integration of long-range input from other regions (Deco et al., 2014, 2017; Sanz-Leon et al., 2015). By determining which optimal parameters best replicate the empirical brain activity of each individual, we can explore systems underlying individual variability in brain activity and common processes shaping network dynamics in health and disease (Klein et al., 2021; Zimmermann et al., 2018).

While global neurobiological characteristics shape individual FC patterns, brain disorders are often related to changes in the properties of only a few areas (Fornito et al., 2015). By exploring which regions play an essential part in maintaining healthy FC, we can discover the neurobiological underpinnings of brain network architecture, as well as potential mechanisms of brain disorders. Previous research suggest that brain networks rely on a few highly connected regions (hub nodes) that significantly affect overall FC (Bullmore & Sporns, 2009; Fransson & Thompson, 2020; Zamani Esfahlani et al., 2020). The strength of these hub nodes is linked to gene expression profiles (Vértes et al., 2016), and their dysfunction is implicated in several brain disorders (Crossley et al., 2014; Royer et al., 2022; Rubinov & Bullmore, 2013). Brain network modelling allows for the precise manipulations of regional neurobiological processes to explore how changes in specific areas affect both static and dynamic FC (Aerts et al., 2016; Alstott et al., 2009).

Traditionally, brain network modelling studies have attempted to replicate the empirical static FC of an individual (Deco et al., 2014; Domhof et al., 2021). While static FC can robustly identify individuals, it cannot capture a wealth of information linked to the temporal evolution of the connectivity pattern (Calhoun et al., 2014; Chiang et al., 2018; Hutchison et al., 2013). Research has shown that dynamic features of FC also vary between individuals (Chen et al., 2016; Davison et al., 2016) and exhibit diagnosis- and symptom-related alterations in brain activity of patients that are not apparent from static FC alone (Pang et al., 2022; White & Calhoun, 2019). These findings suggest that dynamic FC should be considered in investigations of brain alterations.

Here, we present findings from a computational modelling investigation considering dynamic FC. We determined the optimal parameter values that best reproduced empirical static and dynamic FC and the optimal correlation between empirical and simulated static and dynamic FC that could be achieved for each individual. We analysed the associations of fits and parameters with static and dynamic FC features across the brain in order to discover which patterns of FC determine model fits and how neurobiological parameters contribute to individual variability in regional static and dynamic FC. In addition, we investigated the role of different brain regions in generating brain network dynamics by systematically altering the coupling in each region and measuring the resulting changes in static and dynamic FC.

## Methods

### Data Acquisition and Preprocessing

We performed simulations using data from 200 healthy, not related subjects from the Human Connectome Project (HCP) S1200 release (Van Essen et al., 2013). Structural connectivities and parcellated rs-fMRI time courses were generated by Domhof et al. (Domhof et al., 2021, 2022b, 2022a). We used the Desikan-Killiany cortical parcellation (Desikan et al., 2006) for our analyses, which delineates 70 regions of interest based on anatomical structures.

### Computational Modelling

Whole-brain network models describe the brain as a set of regions joined by large-scale connections representing white matter tracts (Sanz-Leon et al., 2015). Each brain area is described as a network of neuronal populations, with a set of parameters regulating the balance between them. Integrating regional activity, the model generates a simulated fMRI signal. In this study, we used the reduced Wong-Wang model with excitatory and inhibitory components (Deco et al., 2014) to simulate the activity in each region. This dynamic mean field model describes the change of the average firing rates and synaptic activities of a population of excitatory and a population of inhibitory neurons with a set of coupled nonlinear differential equations. Model equations are presented in the Supplementary Information. Three forms of input drive the regional activity at each point in time: i) long-range excitatory inputs from other regions, modulated by the structural connectivity, the global coupling parameter G, and the local excitatory synaptic coupling J_N, ii) regional inhibitory currents, modulated by the local feedback inhibitory synaptic coupling J_i, and iii) recurrent excitation, modulated by the local excitatory recurrence w_p and the local excitatory synaptic coupling J_N. We investigated simulated data based on a range of values for G, J_N, J_i, and w_p.

A graphical representation of the modelling approach is shown in Figure 1. We performed all simulations using an average structural connectome, represented by the mean white matter tract weights and lengths across all subjects in the HCP sample. We transformed the two matrices by removing the median and scaling to the interquartile range to obtain a scaling robust to outliers, and rounded to even numbers. We relied on the implementation of the model in the Virtual Brain toolbox (Sanz Leon et al., 2013) to perform the simulations. All simulation settings can be found in Supplementary Table 1.

**Figure 1:**
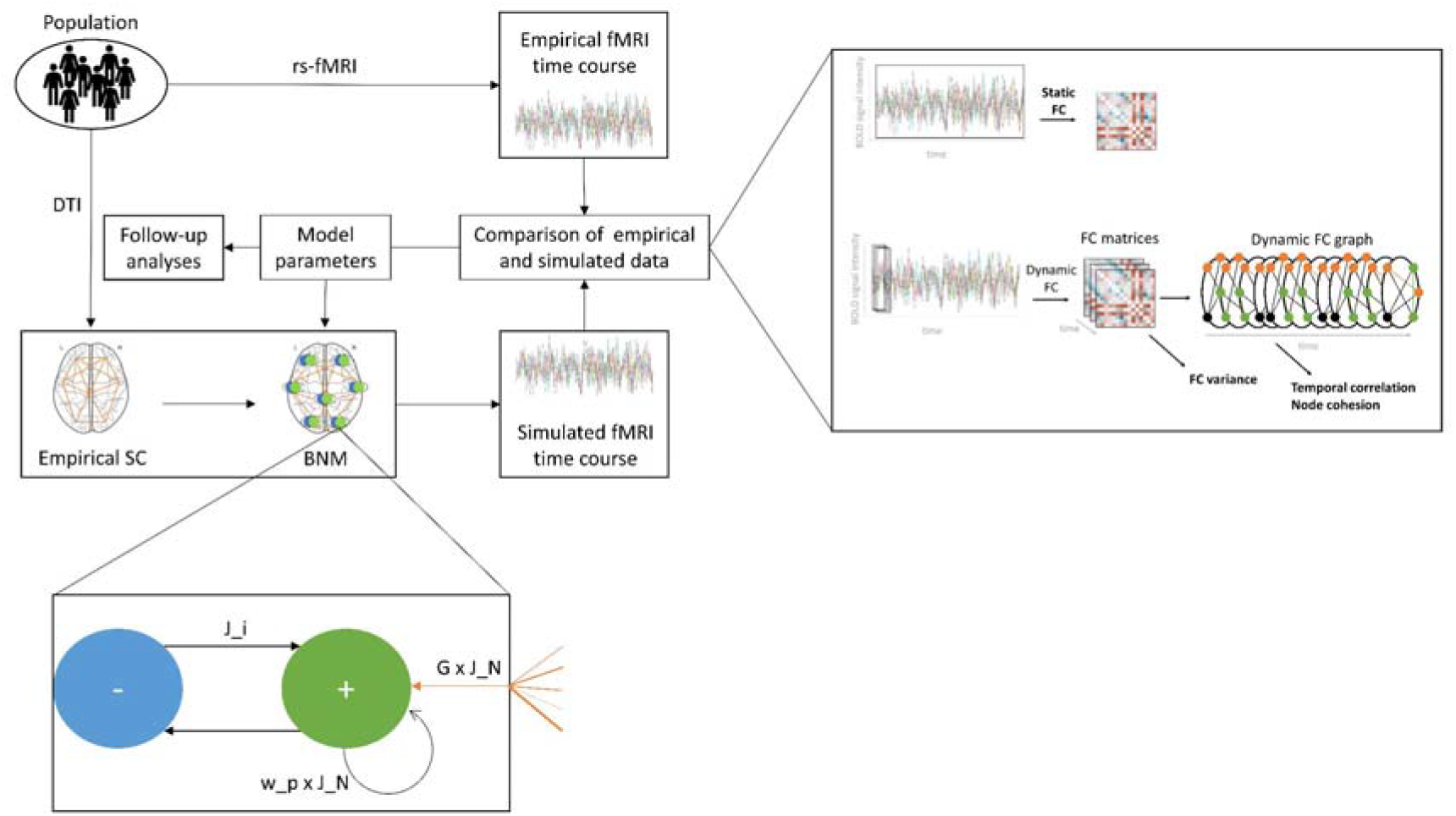
Computational modelling workflow

In order to reduce the parameter space, we performed an initial exploration of parameters to identify a range for J_N, J_i, and w_p at which the model exhibited multistable behaviour for 101 coupling values equally distributed between 0 and 10. In this interval, the model oscillates between two stable states, representing biologically plausible activity. The procedure for this analysis is outlined in the Supplementary Information. We then generated simulations for each combination of 10 values for J_N, J_i, and w_p in the multistability range, amounting to 1000 simulations per coupling value, for a total of 101000 simulations. We obtained a simulated blood oxygen level-dependent (BOLD) fMRI signal in each region by transforming the excitatory synaptic activity using the Balloon-Windkessel hemodynamic model (Buxton et al., 1998; Friston et al., 2003; Mandeville et al., 1999). We then determined the optimal parameter combination by comparing the FC of these simulated time courses with the empirical FC of each subject.

### Calculation of FC measures

We evaluated the models based on both static and dynamic FC. We selected three established metrics which capture important and distinct aspects of dynamic FC (Barber et al., 2021; Sizemore & Bassett, 2018). The static FC (sFC) denotes the correlation in the activity of each pair of regions over the course of the scan, providing a metric for the average of the connectivity over time. FC variance (FCV) is the change in this correlation over time. This measure captures differences in the temporal variability between connections. Temporal correlation (TC) is the consistency of a region’s neighbours from one time point to the next, indicating whether changes in connectivity are abrupt or more gradual. Node cohesion (NC) is the number of times each pair of regions switches communities together. This measure reveals whether nodes are likely to be involved in the same networks and processes. The static FC was determined by calculating the Pearson correlation between the time courses of each pair of regions, yielding a 70×70 matrix for each subject. We used Fisher’s z-transformation to normalise these matrices. In order to compute the dynamic FC parameters, we first calculated the FC in windows of approximately 60 s length, overlapping by 2 s and convolved with a Gaussian kernel of σ = 6 s, producing a 70×70×420 matrix for each subject (Leonardi & Van De Ville, 2015). We then calculated the variance in the connectivity of each pair of regions over time. In addition, we transformed this time course of FC matrices into a dynamic graph by binarising it, keeping only the top 10% of connectivities. From this graph, we first derived the temporal correlation, resulting in a vector of 70 elements. Then, we detected communities of recurrently connected nodes using the tnetwork python library (Cazabet et al., 2021, 2023) and computed the node cohesion, yielding again a 70×70 matrix. The HCP sample contains two resting-state fMRI scans, one recorded using left-right and the other right-left phase encoding, in order to enable researchers to reduce phase encoding-related artefacts by averaging (Van Essen et al., 2013). We computed the dynamic measures separately for each subject, and used the mean of the two matrices or vectors for further analysis.

We calculated the Pearson correlation of these metrics between the simulated and the empirical data. For the measures represented as matrices, we compared only the upper triangulars of the simulated and empirical FC. In order to produce models that reliably reproduced both static and dynamic FC, we identified the model that provided an optimal fit according to both the sFC and the TC using the l2-norm as a global criterion, here referred to as the ‘combined’ metric. In addition, we evaluated the models that produced the highest correlation on each of the individual metrics for comparison. For each of these five optimal models of each subject, we computed the fit across all four of the measures, as well as the corresponding parameter set. We used the optimal model according to the combined metric for all further analyses. We repeated the calculation of optimal parameters and fits and all follow-up analyses in the rs-fMRI data from the second scanning session available in the HCP data set to validate our findings.

### Correlation of fits and parameters with brain-derived features

Based on the optimal models, we considered the relationships between connectivity features and parameters as well as fits. This analysis allows us to discover whether model fits are determined by FC patterns and whether individual variability in FC is related to differences in model parameters. In order to investigate the association between fits and optimal parameters and static and dynamic FC, we computed Pearson correlations of each of the four parameters and the fit according to each of the four measures with a range of features over all 200 subjects of the HCP data. Those features included a) the sFC between each of the 2415 pairs of regions and b) the TC in each of the 70 regions. We performed permutation testing to estimate the significance of the resulting values. A null distribution for each correlation was produced by randomly permuting the parameter or fit variable over the subjects 100000 times and recomputing the correlation. To enhance visualisations, we additionally summarised sFC and TC correlations over seven canonical resting-state networks (Yeo et al., 2011).

To add to this analysis of univariate relationships between model outcomes and individual connectivities, we also attempted to detect multivariate patterns of connectivity features across regions associated with model outcomes. We produced 16 ridge regression models that aimed to predict each parameter or fit value. These models considered either the sFC of the 2415 pairs of regions or the TC in each of the 70 regions as features. For the sFC-based model, we reduced the number of features by performing a principal component analysis (PCA) (Maćkiewicz & Ratajczak, 1993), and selected the minimal number of components that explained more than 90 % of the variance in the data. Additionally, we transformed the smaller number of features for the TC-based model using a PCA, keeping all components. We employed a nested cross-validation (CV) approach to determine generalisation performance (Varma & Simon, 2006). The inner CV cycle determined the optimal alpha value from a range of 100 values logarithmically distributed between 0.01 and 100. In contrast, the outer CV cycle estimated the mean R^2 score of the prediction on previously unseen data. Each level of the CV consisted of 5 folds with 2 permutations, with the PCA transformations estimated individually for each training and test set. In order to determine which regions contributed most strongly to the prediction, we computed the feature importance of each connectivity (Breiman, 2001). We re-trained the model on the entire data set, and calculated the importance of each feature based on the mean drop in the R^2 score over 1000 permutations of that feature. For the sFC-based models, we computed the mean of the importance of the connections of each region to obtain a single importance value for each region. To enhance visualisations, we additionally summarised feature importance over RSNs.

### Region-wise perturbation of parameters

We used a perturbation approach to investigate how FC responds to changes in regional coupling. We systematically altered G in each region and analysed which regional coupling values most strongly impacted FC, and which were particularly important for maintaining biologically accurate FC. We determined a default set of parameters that produce the optimal fit on the combined metric between the simulated data and an sFC matrix and TC vector averaged over the 200 subjects in our sample. We then produced a set of simulations using these default parameters, varying G in each region using 101 values uniformly distributed from 0 to 10. For each of the resulting time series, we computed the sFC, and obtained the global efficiency for each perturbation-derived and the default simulated sFC. The efficiency represents the average inverse shortest path length between each pair of regions, and measures how efficiently information is exchanged across brain networks (Latora & Marchiori, 2001). We determined the cosine distance between the sFC and TC matrices, the difference in the sFC and TC fit of simulated and averaged empirical data, and the difference in the efficiency between the result obtained by simulating with the default parameter values and with the perturbation in each region. In order to determine why certain regions might contribute more strongly to global FC than others, we investigated the relationship between perturbation-related differences and graph properties, dynamic properties and gene expression across the brain. We determined correlations between the mean and variance of the sFC fit difference, TC fit difference, and efficiency difference and three sets of additional features: i) graph metrics including centrality and degree measures of the SC weights and lengths, and the average FC; ii) dynamic metrics of regional BOLD time courses (Lubba et al., 2019) (Supplementary Table 2); and iii) expression data of 15653 genes extracted from the Allen brain atlas (Hawrylycz et al., 2012; Markello et al., 2021). We determined significance levels for each correlation based on nulls generated by permuting the feature importance matrices while preserving their spatial autocorrelation using spin-testing (Alexander-Bloch et al., 2018). These steps were performed with the neuromaps python toolbox (Markello et al., 2022). To facilitate the interpretation of the differential gene expression, we used the MetaScape platform (Zhou et al., 2019) to functionally enrich the genes whose expression correlated significantly (p < 0.05, uncorrected) with the perturbation-induced differences. The gene lists relating to each of the six difference metrics were compared to gene sets included in each Gene Ontology (GO) (GO Consortium, 2004) biological process term, and significantly overrepresented gene sets were extracted. Redundant terms were collapsed into one representative term via clustering.

## Results

While we were able to achieve high fits between empirical and simulated sFC (r = 0.389, σ = 0.037) and TC (r = 0.388, σ = 0.046), the FC variance (r = 0.082, σ = 0.032) and NC (r = 0.196, σ = 0.026) could not be replicated well (Supplementary Figure 1). Using the combined metric of sFC and TC for optimisation, sFC fits averaged 0.339 (σ =0.044) and TC fits 0.378 (σ = 0.044). sFC fit and FCV fit differed significantly between male and female participants (p(sFC) < 0.001, p(FCV) = 0.003), while NC fit differed between the 22-25 and 26-30 age groups (p = 0.039). There were no other significant differences in any fits or parameters between any groups.

The optimal values for the model parameters G and J_N obtained for each subject when considering both static and dynamic FC showed a significant correlation with FC between and TC within several regions (Figure 2i). The sFC of multiple regions showed a strong positive correlation with the coupling parameter G, but a negative correlation with the excitatory synaptic coupling J_N. G exhibited particularly high correlations with connections within the dorsal attention network (DAN), as well as between the somatomotor network (SMN) and the DAN and visual network (VN). For J_N, the strongest associations were present in connections within the VN and those between the frontoparietal network (FPN) and the SMN and VN. The TC correlated strongly positively with G and strongly negatively with J_N across all regions. There were no significant associations between J_i or w_p and sFC or TC in any regions. These findings indicate that G and J_N contribute individual variations in specific static and dynamic FC patterns, while J_i and w_p do not.

**Figure 2:**
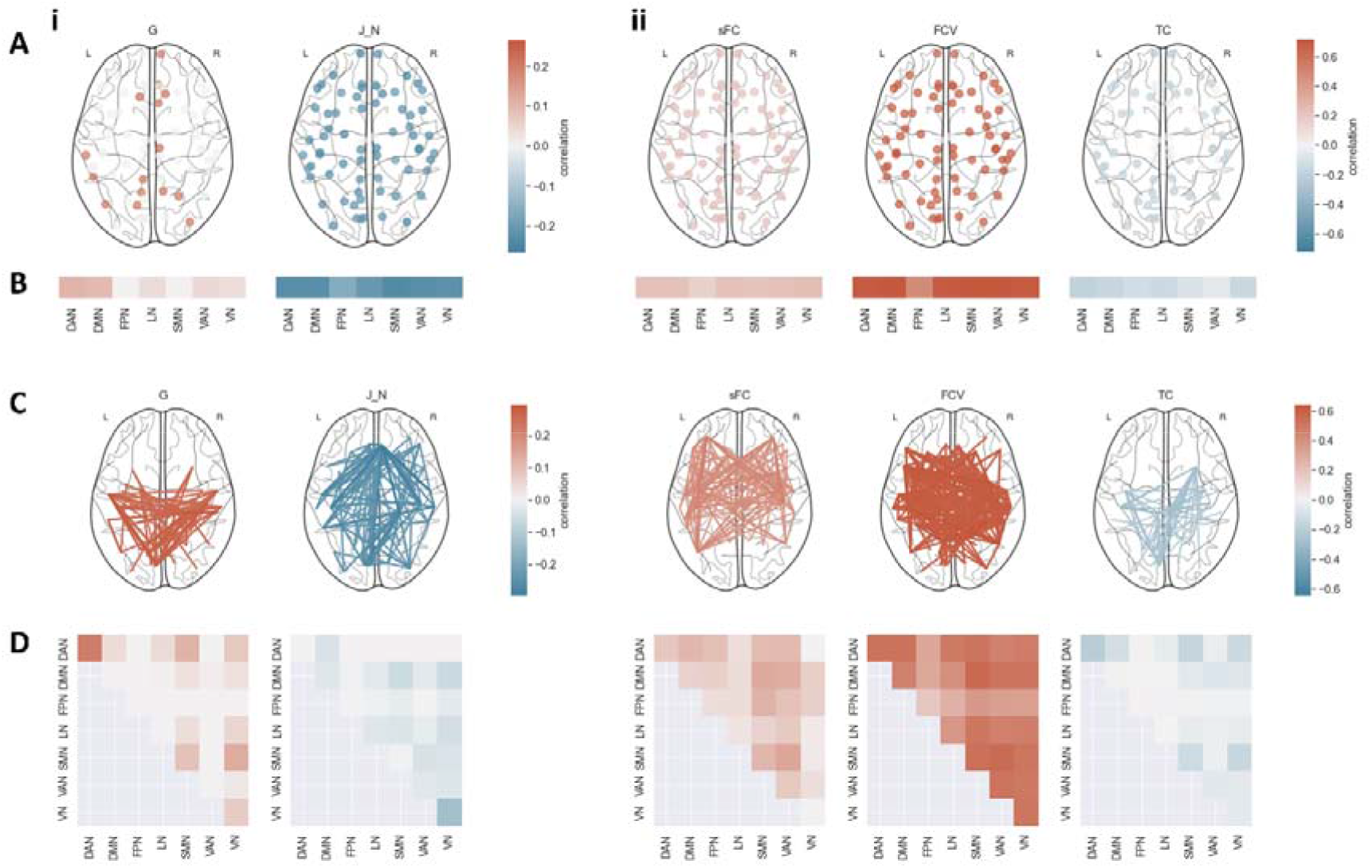
Association between model outcomes and static and dynamic FC features. A Correlation between TC and i) optimal model parameters and ii) optimal fits. Only significant correlations (p_FDR < 0.05) are displayed. B Mean correlation between TC and i) parameters and ii) fits in each resting-state network. Non-significant correlations (p_FDR >= 0.05) were set to 0. C Correlation between sFC and i) parameters, and ii) fits. Only the strongest 30% of significant correlations (p_FDR < 0.05) are displayed. D Mean correlation between sFC and i) parameters and ii) fits within and between resting-state networks. Non-significant correlations (p_FDR >= 0.05) were set to 0. J_i, w_p and NC fit did not exhibit significant correlations with the FC of any region. DAN: dorsal attention network, DMN: default mode network, FPN: frontoparietal network, LN: limbic network, SMN: somatomotor network, VAN: ventral attention network, VN: visual network.

The fit between simulated and empirical data according to the four measures was also strongly connected to some TC and sFC features (Figure 2ii). Subjects with strong sFC also showed high correspondence between empirical and simulated data when considering either sFC or FCV as a target measure. Functional connections between the FPN and the SMN were mainly associated with a strong fit according to sFC. Functional connections between the default mode network (DMN) and the SMN were associated with a strong fit according to FC variance. TC fit exhibited negative correlations with most sFC features, involving the DAN, SMN and VN most strongly. The sFC fit and especially the FC variance fit exhibited strong positive, the TC fit strong negative associations with the TC in most regions. The NC fit did not correlate significantly with sFC or TC in any regions, showing that sFC fit, FC variance fit, and TC fit, but not NC fit, are determined by individual static and dynamic FC patterns.

Multivariate analysis showed that temporal correlation and static connectivity features could predict some model fits reasonably well. Both models predicted the correlation between empirical and simulated FC variance with an R^2^ of above 0.5 (Figure 3A). The fit according to sFC and TC could be predicted with a high score by the model using sFC and TC as features respectively. Neither of the models performed particularly well in predicting the fit according to NC. The model parameters could be best predicted using the TC in each region as features. For J_i and w_p, the achieved scores were close to 0, the R^2 value expected from a constant model which predicts the mean of the target vector for each subject regardless of feature input. The score for G was somewhat below that, while the score for J_N reached a moderate value. The sFC-based prediction model performed worse than a constant model for all parameter targets.

**Figure 3:**
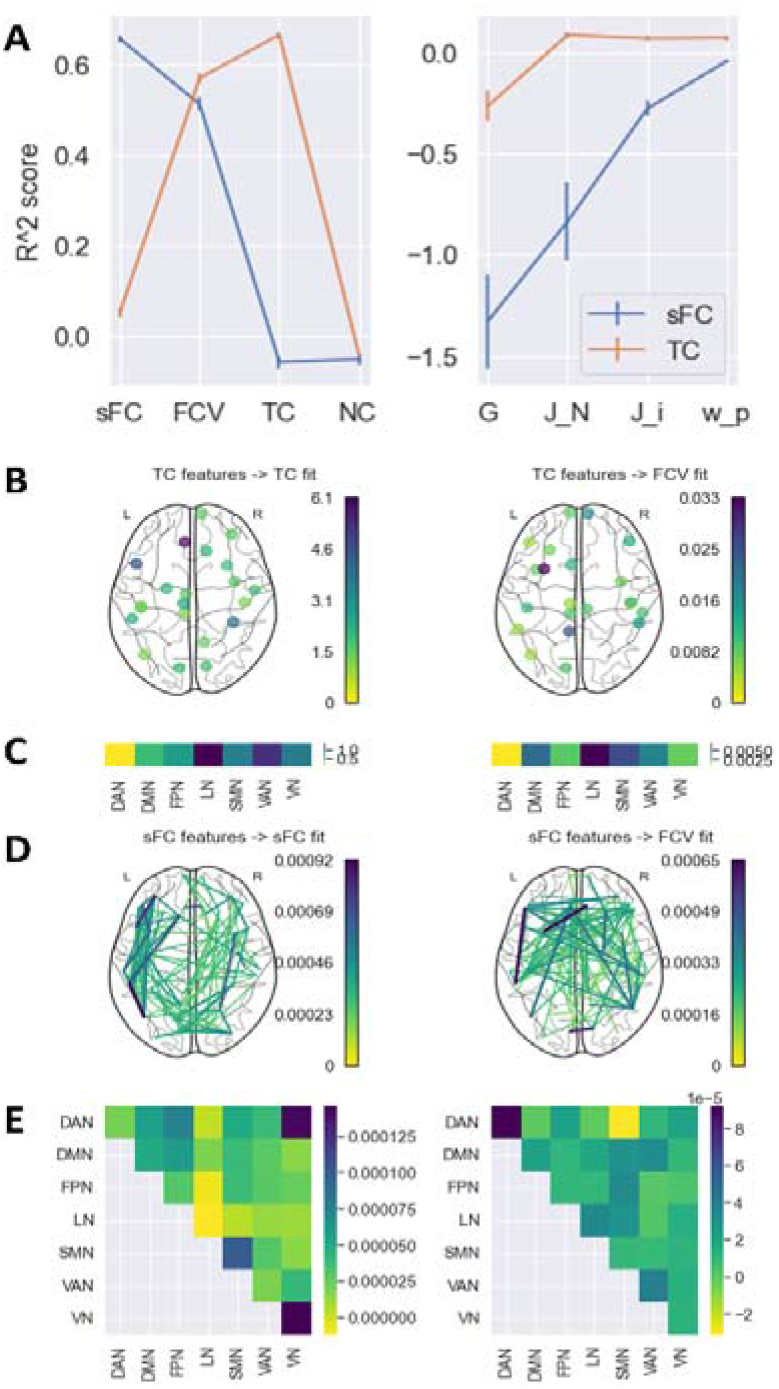
Prediction of fits and parameters from static FC and TC. A Mean R^2 score achieved by models predicting each fit and parameter value based on either the sFC matrix or the TC vector in the outer CV. Error bars represent the standard error of the mean across folds and permutations in the CV. An R^2 of 0 indicates that the model performed as well as a constant model which always predicts the average target value regardless of features. B Feature importance of TC values of each region. Only the top 30% most important regions are displayed. C Mean feature importance of TC values in each RSN. D Feature importance of sFC values of each pair of regions. Only the top 5% most important connections are displayed. E Feature importance of sFC values within and between RSNs. Only predictions with an R^2 score above 0.25 are featured in panels B - E. DAN: dorsal attention network, DMN: default mode network, FPN: frontoparietal network, LN: limbic network, SMN: somatomotor network, VAN: ventral attention network, VN: visual network.

For those prediction models with a high R^2^ score, feature importance analysis showed connectivities that are particularly predictive of the targets (Figure 3B-E). The fit measured by the FC variance was associated most strongly with the TC in the left temporal pole and the left isthmus cingulate, as well as a distributed network of static connections. Regions within the VN, SMN and between the VN and DAN were particularly important for the sFC fit. In contrast, the TC in the LN and VAN, especially in the left medial orbitofrontal gyrus, left pars opercularis, and right fusiform gyrus, had a strong effect on the TC fit.

Perturbation analysis revealed that perturbing the coupling in most regions led to a decrease in fits and efficiency (Figure 4A). However, some regions had a stronger influence on the overall network connectivity than others, while altering the coupling in some regions slightly increased fits and efficiency. The alteration in the efficiency and the sFC fit over all the selected G values was particularly pronounced in the left and right paracentral, the right pre- and postcentral and the right transverse temporal gyrus. The sFC fit could be improved by perturbations in the right medial orbitofrontal gyrus. The effect on the TC fit was much stronger than on the sFC fit, with perturbation in the left pars triangularis in particular causing a reduction, and in the left inferior temporal gyrus an improvement in fit. In some regions, the alterations in fits and efficiency varied strongly depending on the extent of the perturbation. A change in the coupling of the left paracentral gyrus produced a particularly high variance in sFC fit and efficiency differences, while the TC fit varied most strongly when the coupling was perturbed in the left pars triangularis and left pars orbitalis. Over all regions, the fit generally decreased the further G was altered from the benchmark (Supplementary Figure 3).

**Figure 4:**
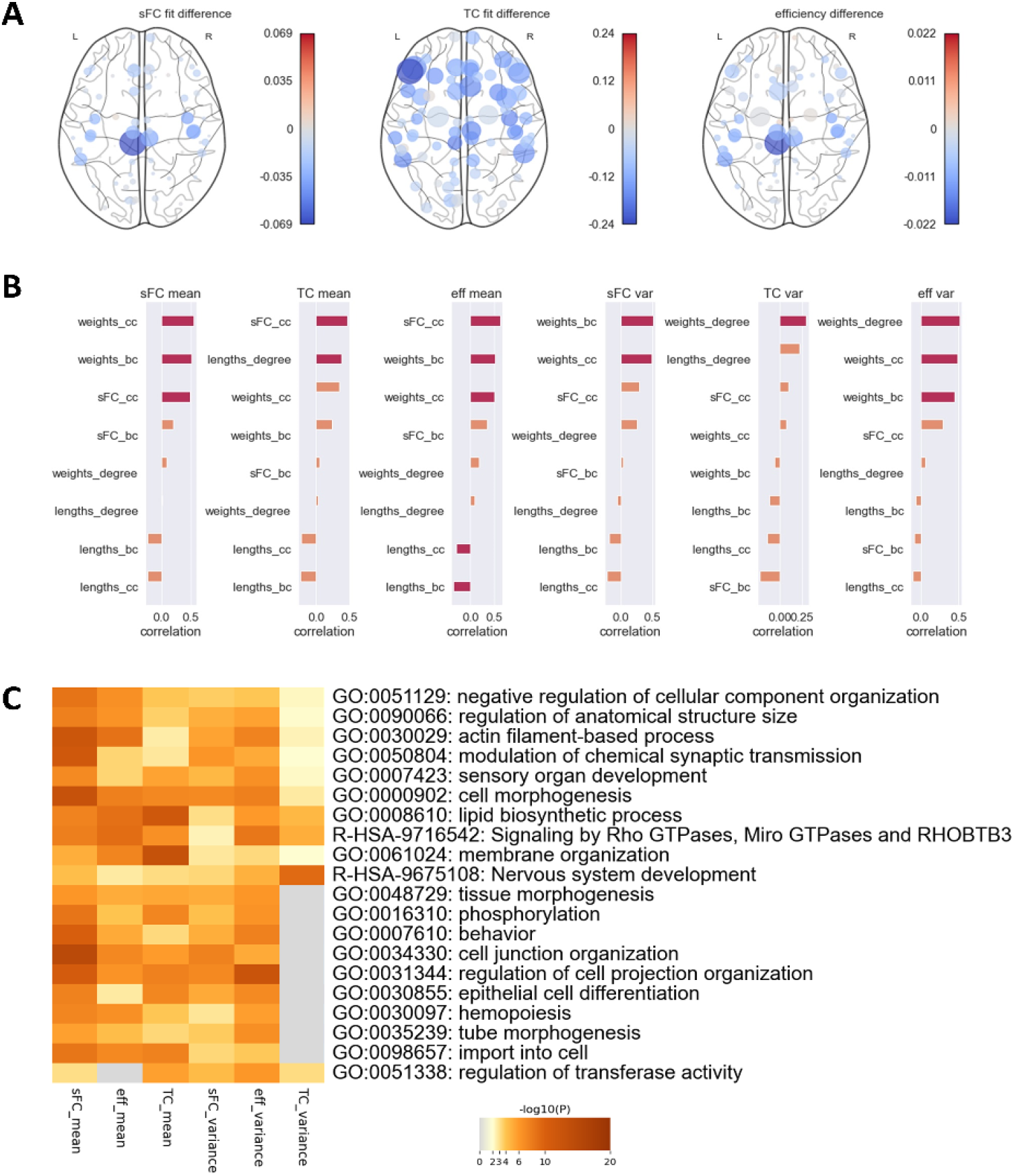
Effects of regional coupling perturbation. A Alterations in i) sFC fit, ii) TC fit, and iii) network efficiency produced by varying the coupling parameter G in each region. Node colors represent the mean alteration. Node sizes represent the variance of the alteration over all G values used. B Correlation of alteration mean and variance of perturbation-induced differences with a range of graph metrics. Orange bars indicate that features did not exhibit significant correlations, red bars show that they correlated significantly before FDR correction (p_uncorrected < 0.05). bc: betweenness centrality, cc: closeness centrality. C Top 10 biological processes involving genes which correlated significantly (p < 0.05) in their expression with mean and variance of perturbation-induced differences.

All six outcome metrics of the perturbation analysis, the mean and variance of the sFC fit difference, the TC fit difference and the efficiency difference, correlated highly with graph parameters of empirical connectivity (Figure 4B). The association with the betweenness centrality and closeness centrality of the SC weights were significant for the means and variances of the sFC fit difference and the efficiency difference. The mean of the TC fit difference correlated most strongly with the closeness centrality of the static FC and the degree of the SC lengths, while the variance correlated most strongly with the degree of the SC weights. Regions which were connected more strongly within the network also led to greater changes in global static and dynamic FC if their coupling was altered. Correlations with dynamic parameters of the regional time courses were weaker and generally did not survive correction for multiple comparisons (Supplementary Figure 4). All six metrics exhibited strong correlations with the expressions of several genes. Enrichment analysis revealed several biological processes involving relevant genes (Figure 4C). Similar pathways were associated with the mean of sFC fit, TC fit and efficiency differences, as well as the variance of sFC fit and efficiency differences, and included brain-related processes such as nervous system development, modulation of chemical synapse transmission, and behaviour. The variance of the TC fit differences was only associated with a subset of the pathways. This structure of association indicates that not all biological processes related to those regional couplings which most impact static FC were associated with the same variability in dynamic FC, but they were associated with regions vital to maintaining biologically accurate global FC.

## Discussion

In this study, we investigated individual neurobiology through brain models which simulate static and dynamic functional connectivity. We then analysed the relationship between optimal model parameters and regional static and dynamic FC features, and determined the effect of each region on overall network connectivity.

We achieved reasonably high maximal fits between simulated and empirical sFC as well as TC for each individual. Given that models optimised for sFC did not manage to reliably replicate empirical TC and vice versa, it was advisable to optimise models for both metrics. When we selected optimal individual models based on a combination of both factors rather than each factor individually, the obtained fits were on par with those reported previously (Klein et al., 2021; Zimmermann et al., 2018). Our models could not replicate empirical NC or FC variance well. The fits of the optimal models were strongly correlated with a set of connectivity features. sFC and FC variance fit showed significant positive correlations with the connectivity in multiple regions, indicating that the models are better able to replicate the sFC of strong and stable connections. This is likely due to the time-independent influence of the underlying structural connectome. TC fit exhibited predominantly negative correlations, showing that the approach favours more random dynamic fluctuations. This effect is presumably caused by the noise added to the regional activity, and suggests that the models can better reproduce arbitrary changes in activity over time, rather than the slow fluctuation between distinct states observed in empirical data (Allen et al., 2014; Vidaurre et al., 2017). sFC features could predict FC variance and sFC fit; in contrast, TC features achieved high prediction scores for FC variance and TC fit, with a small number of features proving to be particularly predictive. These findings show that the modelling framework can better replicate certain connectivity signatures, particularly stronger sFCs within and between the DAN, VAN and LN, and weaker TCs across the brain, particularly in the DAN and VN. Patterns of sFC in SMN, VN and VN-DAN connections, as well as in the TC of LN and VAN, specifically in the left medial orbitofrontal gyrus, left pars opercularis, and right fusiform gyrus, contributed most strongly to fit predictions. The relationship between FC patterns and fit should be considered in future studies, as it provides a potential source of bias if the connectivity in these regions is differentially distributed between groups or inconsistent across sites.

Several of the model parameters exhibited related signatures in the dynamic and static FC. G was generally positively correlated with connectivities, likely because a higher G leads to a higher impact of signals from other areas on regional activity, resulting in increased integration (Sanz-Leon et al., 2015). Connections that were significantly associated with G were concentrated within and between the DAN, SMN and VN. Evidence suggests that multiple neurotransmitters modulate FC in distinct ways. While the impact of dopamine is strongest in the DMN, the VAN (Conio et al., 2020), and the SMN, serotonin affects FC in the DMN, SMN, the FPN and the auditory network (Klaassens et al., 2015). G integrates the effect of disparate systems, suggesting that the regions in which the connectivity correlates with G will see a higher change in connectivity if the overall coupling is altered. Since G scales the combined input from other areas into regional activity, those connections that are strongly linked to G are likely between two regions that receive similar inputs. J_N correlated negatively with the connectivity of regions across the brain, particularly in the VN and in regions connecting the VN and DMN to other networks, although we did not successfully validate these associations in the data from the second scanning session (Supplementary Figure 2). This finding matches previous reports linking disruption in NMDA signalling, via administration of NMDA receptor antagonist ketamine, to brain hyperconnectivity (Anticevic et al., 2015; Driesen et al., 2013) as well as increased dynamic network flexibility (Braun et al., 2016). G and J_N were strongly associated with TC in many regions, indicating that differential moderation of long-range connections and excitatory input is a major driver of TC variability. G correlated positively with regions across the brain, showing that an increase in coupling also leads to increased temporal stability of FC. J_N, on the other hand, correlated negatively with virtually all regions, indicating that increased excitatory signalling might lead to more random FC dynamics.

Perturbation analysis revealed that the coupling in a few regions had a disproportionate effect on overall network connectivity. Altering the coupling in some regions led to particularly large variations in sFC and TC fit and efficiency, suggesting that the coupling level in these regions has an outsized impact on brain connectivity. Other regions exhibited a common shift towards lower fits for most levels of perturbation. These regions appear to be particularly biologically relevant, as the optimal global parameter converged to their regional optimal parameter when fitting the model to empirical data. On the other hand, the few regions in which perturbation led to a mean increase in fit appear to be less relevant to shaping biologically accurate connectivity patterns but could be useful for further improving model accuracy. The fact that the regions which particularly affected sFC fit and efficiency differed from those affecting TC fit indicates that static and dynamic connectivity are governed by distinct neurobiological systems. While the regions central to sFC and efficiency, specifically the left and right paracentral, and the right pre- and postcentral as well as the right transverse temporal gyrus, govern the average functional network architecture, the regions relevant for TC, particularly the left pars triangularis, appear to be vital to the gradual transition between short-term network configurations.

Regions in which perturbation had particularly strong effects exhibited high centrality in the SC, showing that FC is driven to a large extent by regions which facilitate many structural connections. In addition, the means of the perturbation-related differences correlated highly with regional centrality in the functional connectivity, indicating that regions which constitute hubs in the FC are particularly important for maintaining biologically accurate connectivity. These findings support the evidence from lesion modelling studies, which suggests that disturbances in hub nodes (Achard et al., 2006; Aerts et al., 2016), particularly those along the cortical midline (Alstott et al., 2009), have the strongest effect on FC. The role of the regions we identified, particularly the pre-, post- and paracentral gyri, in maintaining normal brain connectivity further explains the observation that these regions are particularly well protected against stroke (Thirugnanachandran et al., 2024).

The expression of genes associated with a number of biological processes was related to the strength of the perturbation effect. The relevant mechanisms included some specific to the central nervous system, such as nervous system development and behaviour. Other, more general processes, such as membrane organisation and phospholipid metabolism, might nonetheless influence brain connectivity, as they are involved in myelination (Raasakka et al., 2017), a key factor in preserving connections between brain regions (Huntenburg et al., 2017; Vandewouw et al., 2021). In addition, the impact of regional perturbations also correlated strongly with the expression of genes relating to the modulation of chemical synapse signalling, indicating that regions which affect FC most strongly exhibit more neuromodulatory activity. This further supports the hypothesis that neuromodulation and neurotransmission are essential contributors to the establishment and maintenance of global FC.

While the majority of our findings could be replicated in a second set of resting-state fMRI data of the same participants (Supplementary Figure 2), some limitations should be considered in the interpretation of these results. Firstly, we modelled individual brain activity based on an averaged structural connectome for all subjects. While this provided a clearer insight into the relationship between FC and model parameters, employing individual connectomes could help elucidate individual structure-function relationships. Further, fitting model parameters individually for each region could contribute to an even more detailed picture of individual neurobiology. Given that we found only a subset of regional connectivity features reflected individual variability in model parameters, analysing these relationships in patients with brain disorders might provide new information on brain alterations.

In this study, we showed that modelling dynamic FC allows us to investigate neurobiological correlates of brain network dynamics. We found that differences in long-range inputs drive individual variability in some aspects of global dynamic FC, while variability in static FC is shaped by a combination of regional parameters. In addition, we identified that the coupling in a subset of frontal regions has a major impact on global network connectivity. A future investigation of the relationships between neurobiology and dynamic FC in brain disorders might reveal new insights into the origins of brain abnormalities in patients.

## Author Contributions

Linnea Hoheisel: Conceptualisation, Methodology, Software, Formal Analysis, Visualisation, Writing - Original Draft, Writing - Review & Editing. Hannah Hacker: Software, Writing - Review & Editing. Gereon R Fink: Writing - Review & Editing. Silvia Daun: Supervision. Joseph Kambeitz: Conceptualisation, Methodology, Supervision, Writing - Review & Editing.

## Acknowledgements

Data were provided by the Human Connectome Project, WU-Minn Consortium (Principal Investigators: David Van Essen and Kamil Ugurbil; 1U54MH091657) funded by the 16 NIH Institutes and Centers that support the NIH Blueprint for Neuroscience Research; and by the McDonnell Center for Systems Neuroscience at Washington University.

