## Supplementary Material for "Computational modelling reveals neurobiological contributions to static and dynamic functional connectivity patterns"

**Computational Modelling of Brain Network Dynamics reveals neurobiological contributions to static and dynamic functional connectivity patterns**

Linnea Hoheisel^1,2^, Hannah Hacker^2^, Gereon R Fink^1,3^, Silvia Daun^1,4^, Joseph Kambeitz^1,2^

*1. Institute of Neuroscience and Medicine (INM-3), Forschungszentrum Jülich, Jülich, Germany*

*2. Department of Psychiatry and Psychotherapy, University Hospital of Cologne and Faculty of Medicine, University of Cologne, Cologne, Germany*

*3. Department of Neurology, University Hospital of Cologne and Faculty of Medicine, University of Cologne, Cologne, Germany*

*4. Institute for Zoology, University of Cologne, Cologne, Germany*

Model equations

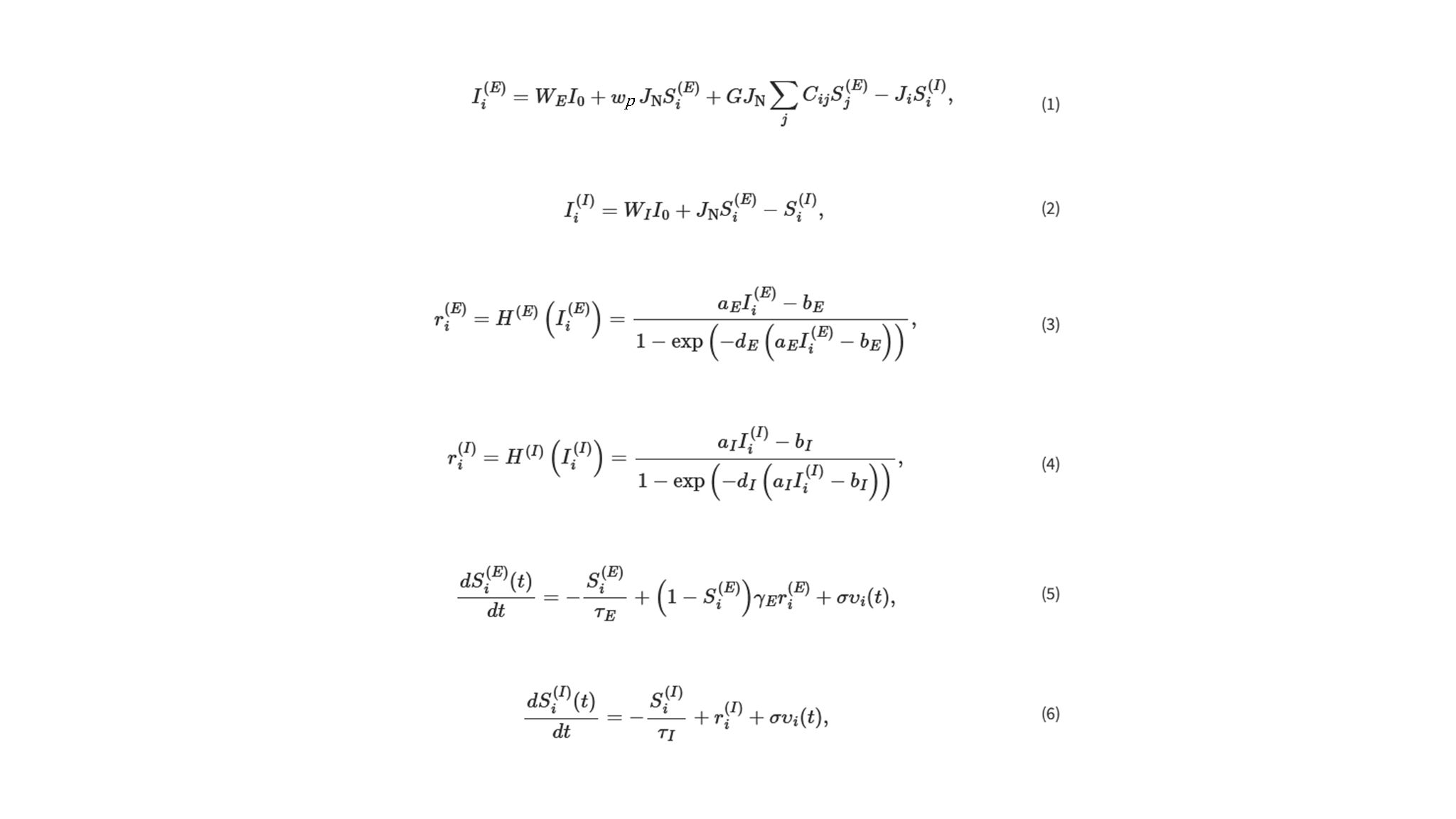

*r_i_^(E,I)^*: firing rate in excitatory *(E)* or inhibitory *(I)* population of node *i*; *S_i_^(E,I)^* average synaptic gating variable; *I_i_^(E,I)^* : input current; *G*: global coupling ; (*C_ij_* ): structural connectivity between each pair of regions *i* and *j*; *J_i_*: local feedback inhibitory synaptic coupling; *I_0_*: external input; *w_(E,I)_*: population external input scaling weight; *w_p_*: local excitatory recurrence; *J_N_*: excitatory synaptic coupling

Identification of bifurcation range

In the bifurcation interval, the model can reach two stable states, and converges to a high or a low state in the absence of noise depending on the initial conditions. If noise is added to the model, a biologically plausible oscillation is generated in this range. For a given value for G, we generated deterministic simulations of 10 seconds with initial conditions representing high or low activity in all regions. We tested each combination of five values for J_N, J_i and w_p distributed evenly in the range suggested by Deco et al. (Deco et al., 2014), amounting to 250 simulations in total. We obtained a time course of the state variable S_i, the average synaptic gating variable in each region, and determined the maximum value at the last simulated time point for both the low and high initial conditions.

We then calculated the difference between the maximal S_i for the high and low conditions for each set of parameters. If this yielded parameter combinations for which the difference was higher than 0.2, indicating multistability, we selected these combinations to test for the optimal parameter range. If we could not identify such a combination of parameters directly, we iteratively increased the resolution of parameters to test. We determined the point at which the maximal S_i for the low initial condition showed the greatest change, indicating a switch from a system with a low attractor state to one with a high attractor state, along the three axes representing the parameters J_N, J_i and w_p. The system becomes multistable as it transitions between the low-attractor and the high-attractor regimes. Thus, in the next iteration, we considered five values equally distributed in the range in which the largest difference in the maximal S_i for the low initial condition occurred for each of the three parameters. We repeated this procedure until we could identify a range of parameters for which the difference between the maximal S_i for the high and low conditions exceeded 0.2, representing the multistability range, and used this range as a set of biologically plausible parameters for this value of the coupling parameter G.

Supplementary Tables

| stage | monitor | period | integrator | integration step | length | I_0 | G | J_N | J_i | w_p |
| --- | --- | --- | --- | --- | --- | --- | --- | --- | --- | --- |
| parameter exploration | TemporalAverage | 1 | EulerDeterministic | 0.2 | 10^4 ms | 0.3 | start = 0  stop = 10  step = 0.1 | start = 0.001  stop = 0.5  step = 0.1 | start = 0.001  stop = 2.0  step = 0.4 | start = 0  stop = 2  step = 0.4 |
| simulation | Bold | 2000 | EulerStochastic | 1 | 10^6 ms | 0.3 | start = 0  stop = 10  step = 0.1 | dependent on G | dependent on G | dependent on G |
| perturbation | Bold | 2000 | EulerStochastic | 1 | 10^6 ms | 0.3 | 1.4 | 0.5 | 2.0 | 0.67 |

**Supplementary Table 1: Simulation settings**

| **Feature name** | **Definition** |
| --- | --- |
| DN_HistogramMode_5 | Mode of *z*-scored distribution (5-bin histogram) |
| DN_HistogramMode_10 | Mode of *z*-scored distribution (10-bin histogram) |
| SB_BinaryStats_mean_longstretch1 | Longest period of consecutive values above the mean |
| DN_OutlierInclude_p_001_mdrmd | Time intervals between successive extreme events above the mean |
| DN_OutlierInclude_n_001_mdrmd | Time intervals between successive extreme events below the mean |
| CO_f1ecac | First 1 / *e* crossing of autocorrelation function |
| CO_FirstMin_ac | First minimum of autocorrelation function |
| SP_Summaries_welch_rect_area_5_1 | Total power in lowest fifth of frequencies in the Fourier power spectrum |
| SP_Summaries_welch_rect_centroid | Centroid of the Fourier power spectrum |
| FC_LocalSimple_mean3_stderr | Mean error from a rolling 3-sample mean forecasting |
| CO_trev_1_num | Time-reversibility statistic |
| CO_HistogramAMI_even_2_5 | Automutual information, m = 2, τ = 5 |
| IN_AutoMutualInfoStats_40_gaussian_fmmi | First minimum of the automutual information function |
| MD_hrv_classic_pnn40 | Proportion of successive differences exceeding 0.04 σ (Mietus et al., 2002) |
| SB_BinaryStats_diff_longstretch0 | Longest period of successive incremental decreases |
| SB_MotifThree_quantile_hh | Shannon entropy of two successive letters in equiprobable 3-letter symbolization |
| FC_LocalSimple_mean1_tauresrat | Change in correlation length after iterative differencing |
| CO_Embed2_Dist_tau_d_expfit_meandiff | Exponential fit to successive distances in 2-d embedding space |
| SC_FluctAnal_2_dfa_50_1_2_logi_prop_r1 | Proportion of slower timescale fluctuations that scale with DFA (50% sampling) |
| SC_FluctAnal_2_rsrangefit_50_1_logi_prop_r1 | Proportion of slower timescale fluctuations that scale with linearly rescaled range fits |
| SB_TransitionMatrix_3ac_sumdiagcov | Trace of covariance of transition matrix between symbols in 3-letter alphabet |
| PD_PeriodicityWang_th0_01 | Periodicity measure of (X. Wang et al., 2007) |

**Supplementary Table 2: 24 dynamic parameters of regional time courses (Lubba et al., 2019) considered in the correlation analysis and perturbation enrichment analysis.**

Supplementary Figures

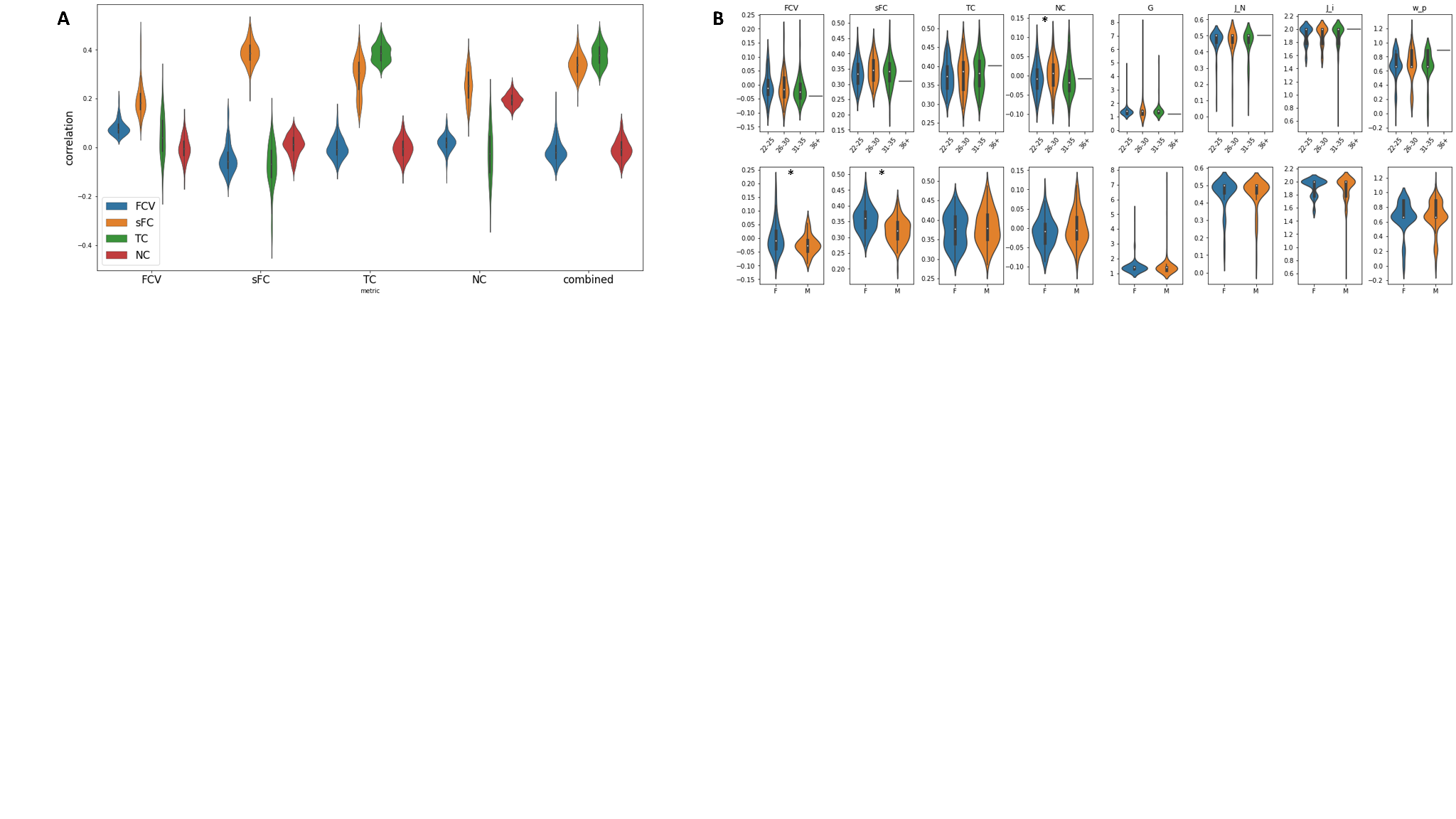

**Supplementary Figure 1: Simulation outcomes**

**A Correlations between simulated and empirical data of optimal solutions based on each of the five metrics considered. When models are optimised for the sFC or the TC, they score highly on those measures, but far lower on each of the other three measures. Models optimised for the combined metric of sFC and TC score well on both of those measures, but show substantial individual variability in the fit. The simulations cannot replicate the FCV or the NC well, with even the models optimised for those metrics only achieving low correlations. B Parameters and fits split by covariate values. Stars indicate significant differences (p < 0.05).**

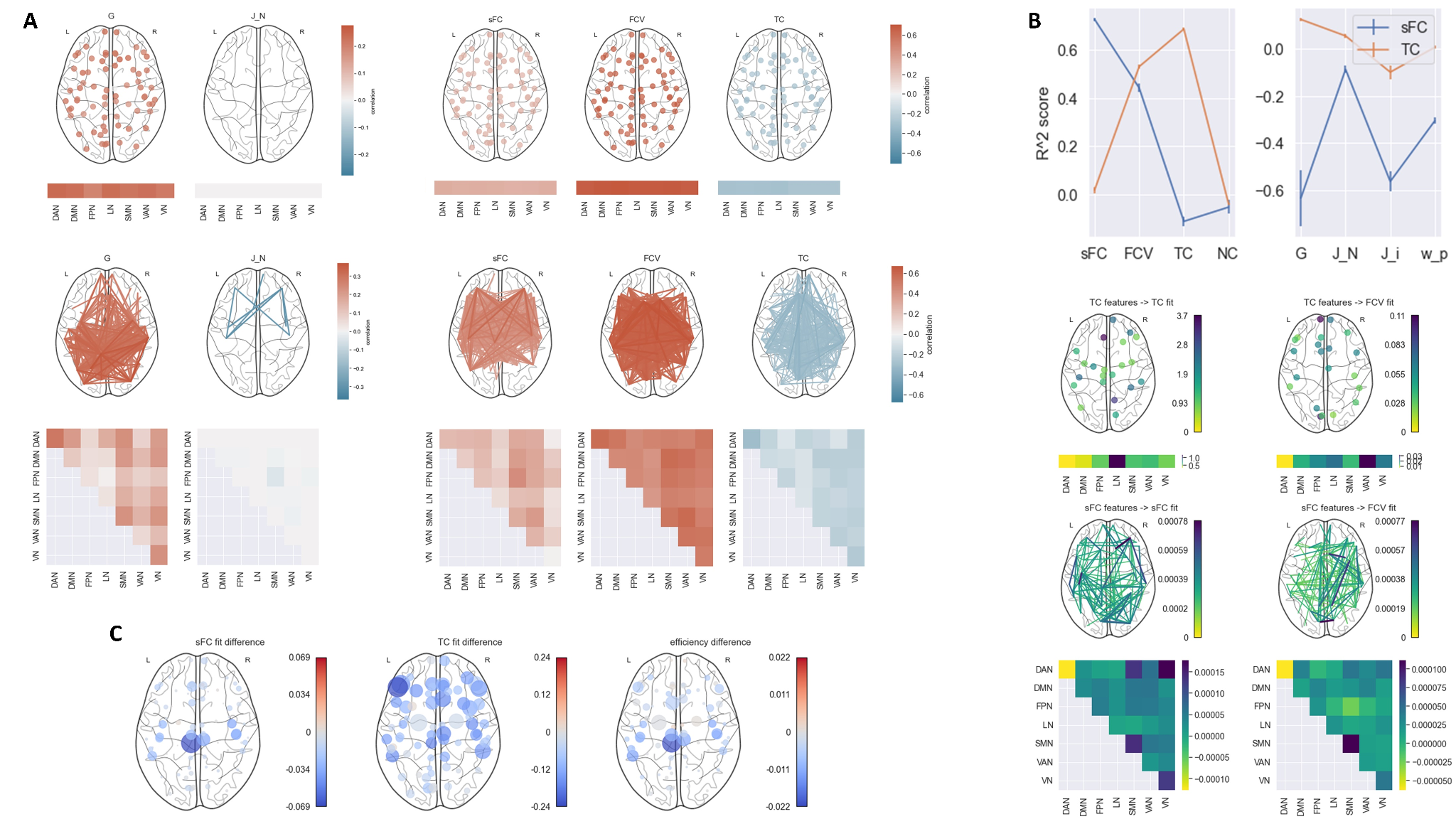

**Supplementary Figure 2: Validation of results.**

**A Results of correlation analysis. B Results of machine learning analysis. C Results of perturbation analysis. The results largely matched those of our original analysis. The FC features correlated with G, sFC fit and FCV fit remained largely the same in both sessions. In the first session, we identified more significant FC correlations of J_N, but fewer of TC fit, and fewer TC correlations of G. We were also able to replicate the machine learning results, with the same models achieving similar scores. The feature importance also remained largely the same across the two sessions, although the importance of sFC features for the prediction of FCV fit was altered somewhat. Perturbation analysis revealed the same relevant regions in both the first and the second session.**

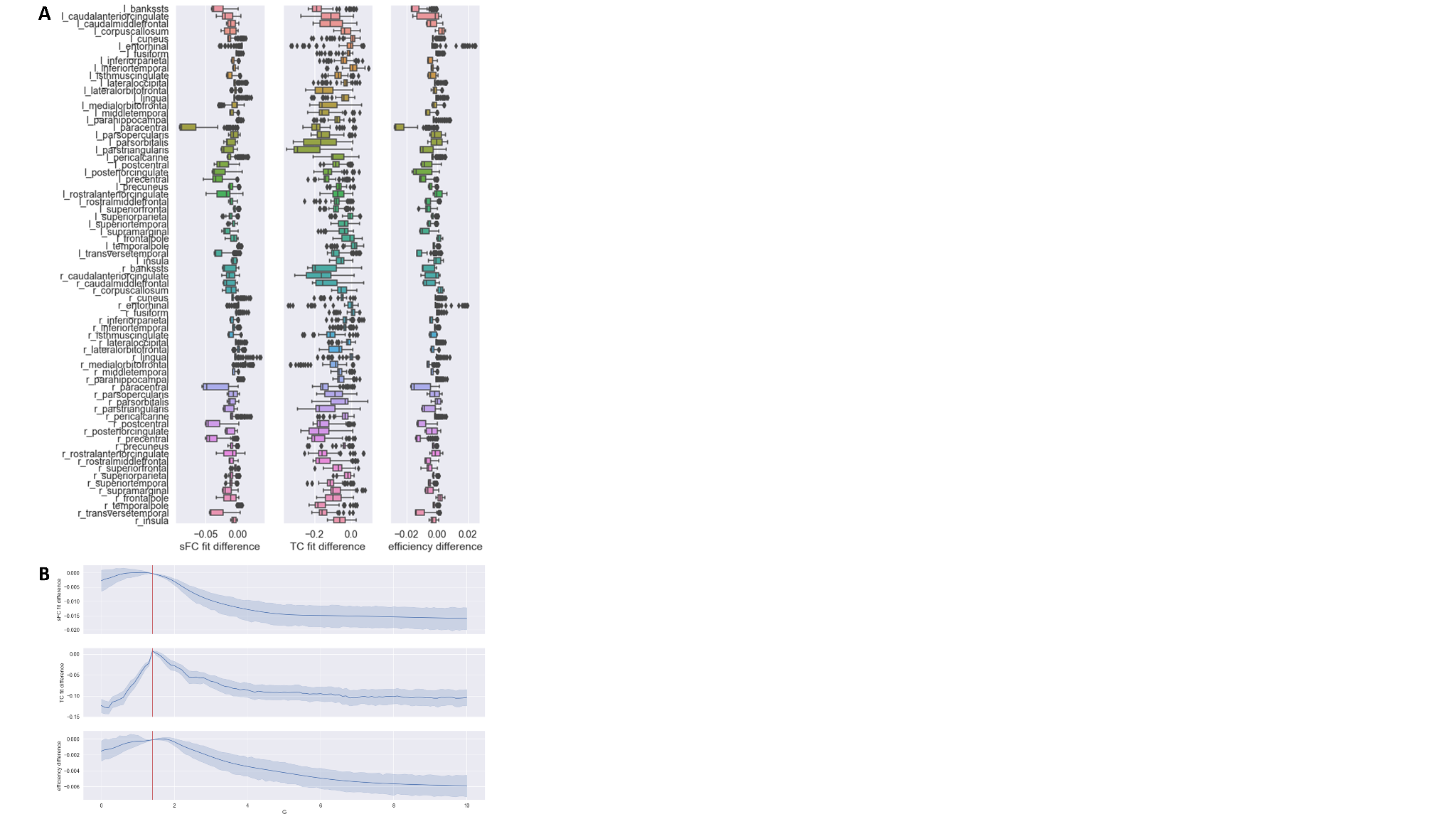

**Supplementary Figure 3: Perturbation of coupling in each region**

**A Difference in sFC fit, TC fit, and efficiency between perturbation-derived and default simulations for each perturbed region.**

**B Difference in sFC fit, TC fit, and efficiency between perturbation-derived and default simulations as a function of G in the perturbed region. Red lines indicate the default value.**

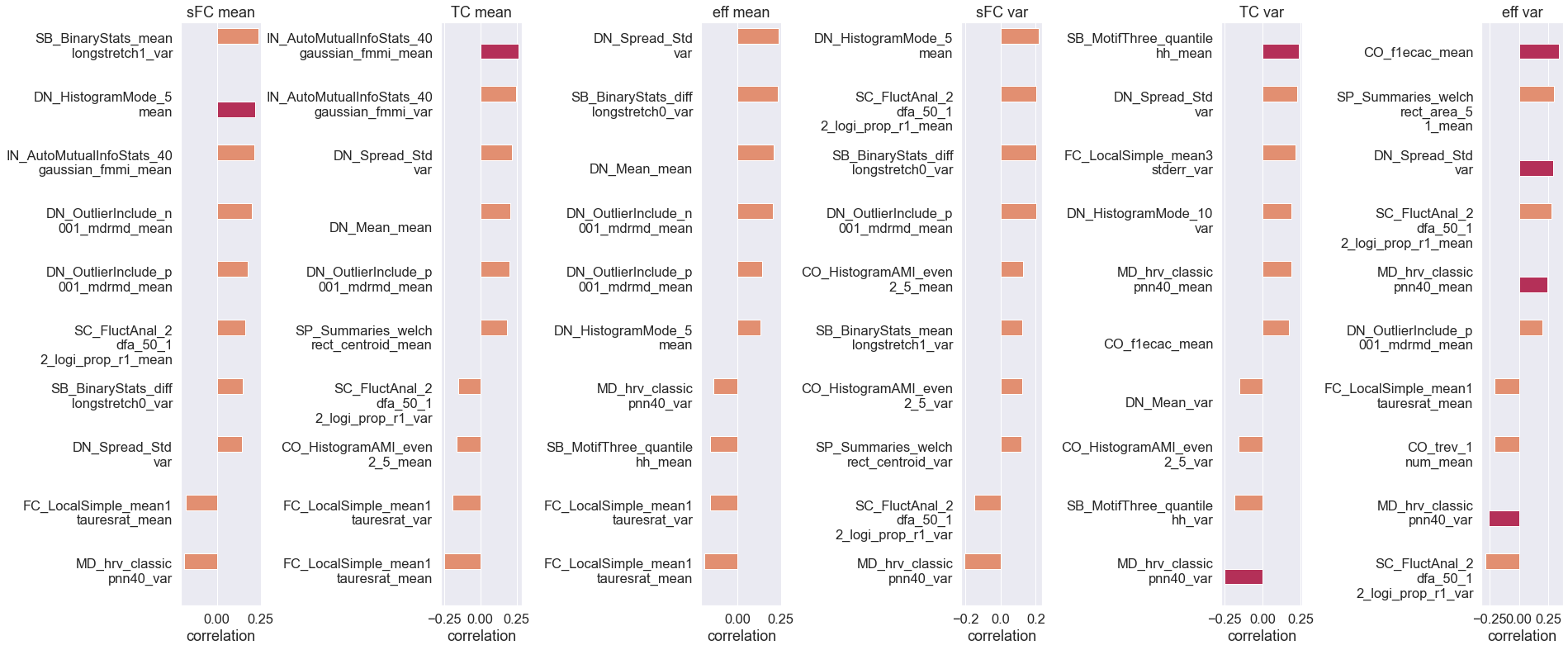

**Supplementary Figure 4: Correlation of alteration mean and variance of perturbation-induced differences with a range of graph metrics.**

**Orange bars indicate that features did not exhibit significant correlations, red bars that they correlated significantly before FDR correction (p_uncorrected < 0.05).**
